# DeepCOMBI: Explainable artificial intelligence for the analysis and discovery in genome-wide association studies

**DOI:** 10.1101/2020.11.06.371542

**Authors:** Bettina Mieth, Alexandre Rozier, Juan Antonio Rodriguez, Marina M.-C. Höhne, Nico Görnitz, Klaus-Robert Müller

## Abstract

Deep learning algorithms have revolutionized data science in many fields by greatly improving prediction performances in comparison to conventional approaches. Recently, explainable artificial intelligence (XAI) has emerged as a novel area of research that goes beyond pure prediction improvement. Knowledge embodied in deep learning methodologies is extracted by interpreting their results. We investigate such explanations to explore the genetic architectures of phenotypes in genome-wide association studies. Instead of testing each position in the genome individually, the novel three-step algorithm, called DeepCOMBI, first trains a neural network for the classification of subjects into their respective phenotypes. Second, it explains the classifiers’ decisions by applying layerwise relevance propagation as one example from the pool of XAI techniques. The resulting importance scores are eventually used to determine a subset of most relevant locations for multiple hypothesis testing in the third step. The performance of DeepCOMBI in terms of power and precision is investigated on generated datasets and a 2007 WTCCC study. Verification of the latter is achieved by validating all findings with independent studies published up until 2020. DeepCOMBI is shown to outperform ordinary raw p-value thresholding as well as other baseline methods. Moreover, two novel disease associations (rs10889923 for hypertension and rs4769283 for type 1 diabetes) were identified.

## Introduction

Genome-wide association studies (GWAS) investigate the phenotypic effects of small genetic variations called Single Nucleotide Polymorphism (SNPs). While some methods for the analysis of GWAS focus on phenotypic risk prediction based on the given genetic information^1,2,3,4,5^, others try to explain these risk effects by highlighting which SNPs are having an effect on a given trait^6,7,8,9,10^. This work aims at a combination of both of these goals and uses a deep learning based prediction method in combination with statistical testing to identify SNPs associated with the phenotype under investigation.

Following developments in biotechnology, the first GWAS was published in 2002^11,12,13^. Several years later, a landmark study - the largest GWAS ever conducted at the time of its publication in 2007 - was presented by the Wellcome Trust Case Control Consortium (WTCCC)^14^ including 14,000 cases of seven common diseases and 3,000 shared controls. Ever since then, sample sizes, rates of discovery and numbers of traits studied have been rising continuously^15^. According to the GWAS catalog accessed on September 15th 2020 >4,700 studies have investigated more than 3,500 phenotypes and identified >200,000 SNP phenotype associations with *p*-values below 1*10^−5^. Especially for common human diseases such as diabetes, autoimmune disorders or psychiatric illnesses GWAS have provided valuable insight into the corresponding genetic inheritance processes^16^. A few studies have included over 1 million subjects enabling the identification of SNPs with lower risks and frequencies^17,18^.

However, the vast amount of available data on SNP phenotype associations still only accounts for a small fraction of heritability. The genetic architectures and variances of most traits and diseases remain largely unexplained. This effect, often referred to as “the missing heritability”, is assumed to - at least partially - be caused by the way GWAS datasets are traditionally analyzed^19,20^. The classic approach - which we refer to as raw *p*-value thresholding (RPVT) - consists of carrying out a statistical association test to assign a *p*-value to each SNP under investigation and subsequently assessing its statistical significance via a comparison to a predefined threshold *t\**.^16^ This standard approach to analysing GWAS is therefore based on testing SNPs individually and in parallel, which intrinsically ignores any potential interactions^20,21^ between or correlation structures among the set of SNPs under investigation^22,23,^. Studies fail to identify multi-locus effects by using the traditional RPVT approaches and a large amount of potentially available information is lost^24^. Only very few diseases rely on single genetic defects with large effects. Most complex diseases are caused by epistatic interactions of multiple genetic factors with small effect sizes, which are further influenced by correlation structures due to both population genetics and biological relations^15^. Brute force multivariate approaches to identify such dependencies are oftentimes computationally too expensive for large GWAS datasets and are limited by low statistical power due to excessive multiple testing. A few attempts have been made to identify genetic interactions, but most of them were not able to find strong, statistically significant associations^21,25,26,27^.

To overcome these limitations of traditional approaches and following the rise of machine learning in data science and an increasing amount of available large-scale GWAS datasets, a number of methods have been proposed to introduce machine learning tools for the analysis of such studies. Linear approaches such as multivariate logistic regression and sparse penalized methods including Lasso have been applied to GWAS datasets. In general, penalized models achieve better performances than non-penalized methods^4,28,29,30^. Some of the top-performing models combine statistical testing and machine learning for the identification of SNP disease associations^6,7,31^. While most of these methods do not provide validation on real data comparing to the GWAS database, very few provide a full evaluation of identified genetic variants in terms of comparison to previously published GWAS. Other proposed nonlinear models, such as random forests, gradient boosted trees and bayesian models^4,28,32,33^ investigate interactions and correlations in the genetic architecture of traits, but were mostly found to be outperformed by linear penalized methods^4,25,28,33^.

To harness even more sophisticated nonlinear machine learning methods for the analysis of GWAS, attention has recently been drawn to deep neural networks (DNN). This powerful tool for learning nonlinear relationships between an input and an output variable by transferring information through *“a computing system made up of a number of simple, highly interconnected processing elements”*^34^ has seen an unprecedented rise in data science^35^ and created enormous progress in numerous fields, e.g. image classification^36,37^, natural language processing^38^, speech recognition^39^ and quantum chemistry^40^. DNNs have been applied to the analysis of GWAS datasets^41,42^, but most of the corresponding publications focus on risk prediction^28,43,44,45^ and only very few methods have been proposed for the identification of SNP disease associations^28,46^.

Romagnoni et al.^28^ present a thorough comparison of conventional statistical approaches, traditional machine learning based techniques and state-of-the-art deep learning based methods in terms of both prediction rates and the identification of SNP associations on a Crohn’s Disease immunochip dataset. Classification performances of numerous methods (Lasso as reference, penalized logistic regression, gradient boosted trees, DNNs) were compared and found to be similar for most methods (linear and nonlinear) implicating potentially “limited epistatic effects in the genetic architecture”^28^. However, when investigating the associated genetic regions identified by the different methods, machine learning and deep learning based methods were indeed found to provide new insights into the genetic architecture of the trait. Romagnoni et al. achieved this by applying the concept of explainable AI, which is an emerging field of AI that has been gaining importance recently^47^. It refers to techniques, which open the so-called “black box” of machine learning methods and reveal the processes underlying their decisions so that the results can be better understood. The explanation method used by Romagnoni et al. - permutation feature importance (PFI) - is a generalized, model-agnostic approach and more sophisticated methods specifically tailored to DNNs are available. To the best of our knowledge, deep Taylor based explanation techniques^48^ have not yet been applied in the field of GWAS and we propose to adopt layerwise relevance propagation (LRP)^49,50^ for the analysis of such data. LRP is a direct way to compute feature importance scores and has been applied very successfully in numerous data science problems to explain decisions of DNNs^51,52^. Instead of basing the importance score of a SNP on the data of that SNP alone, correlation structures and possible interactions are automatically taken into account.

To make LRP applicable as an explanation method for GWAS data, we use a very promising, well performing machine learning based method, called COMBI^31^, as a starting point for our deep learning based approach. COMBI is a two-step method, which first calculates a relevance score for each SNP by training a support vector machine (SVM)^53^ for the classification of subjects based on their genetic profile. Using the learned SVM weights as an indicator of importance, COMBI selects the highest scoring SNPs as a subset to put into multiple hypothesis testing. This approach was shown to outperform other combinatorial approaches and a number of purely statistical analysis tools. The method we propose here can be viewed as an extension of the COMBI method^31^ replacing the rather simple prediction step of an SVM with a more sophisticated deep learning method and using the concept of explainability to extract SNP relevance scores via LRP.

We propose a deep learning based approach for the identification of SNP phenotype associations and call the novel method DeepCOMBI (See **Figure 1**). The three step algorithm consists of

1. a deep learning step where we train a DNN for classifying individuals based on their SNP data;
2. an explanation step where we calculate SNP relevance scores by applying LRP and reduce the number of SNPs by selecting only the most explanatory SNPs; and
3. a statistical testing step where only the SNPs selected in step 2 are tested for statistically significant association with the trait under investigation.

**Figure 1.**
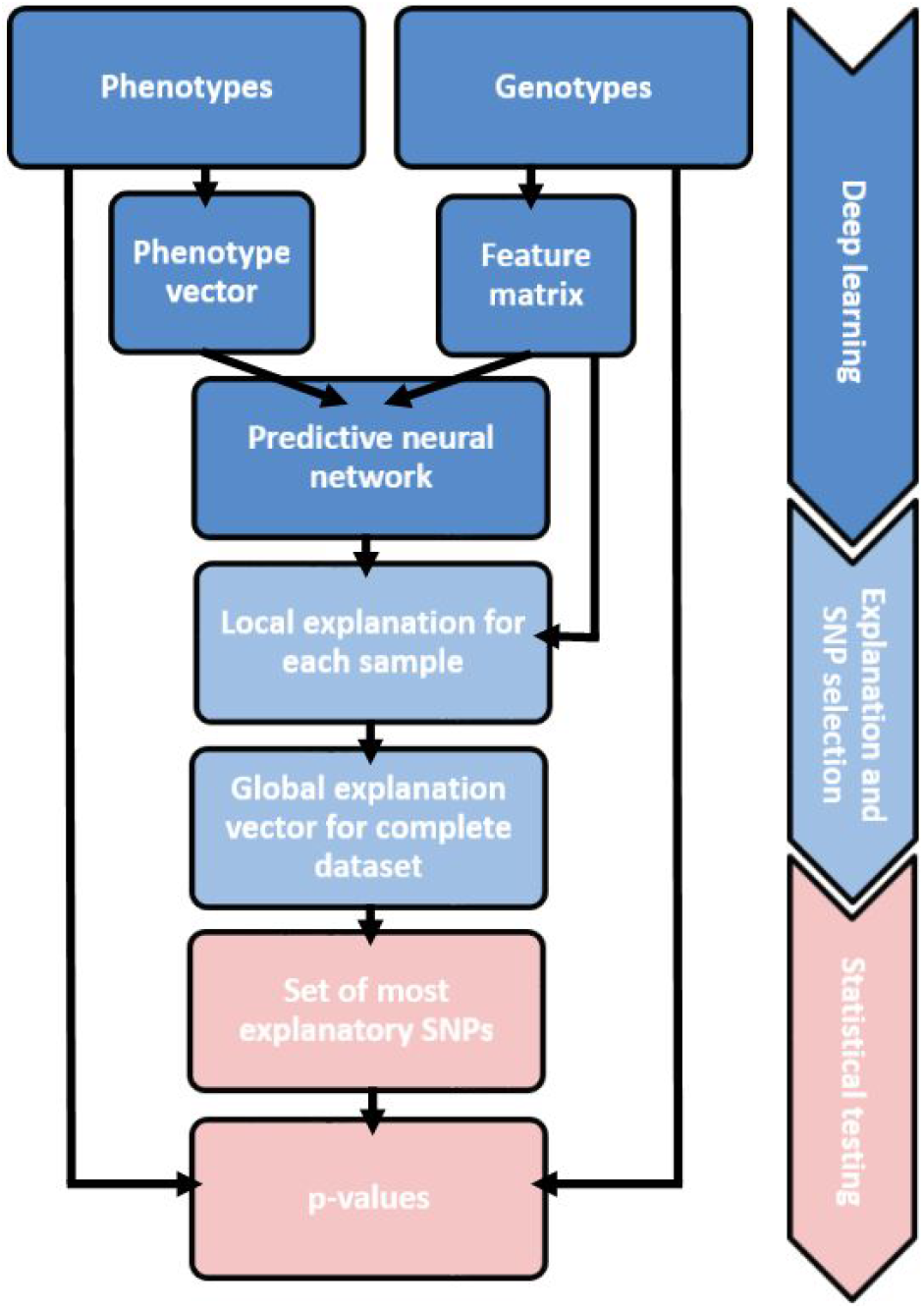
Overview of the DeepCOMBI method. Receiving genotypes and phenotypes of a GWAS as input, the DeepCOMBI method first applies a deep learning step to train a DNN for the classification of subjects. Afterwards, in the explanation step, it selects the most relevant SNPs by applying LRP to calculate relevance scores for each SNP. Finally, for this set of most relevant SNPs, DeepCOMBI calculates *p*-values and corresponding significance thresholds in a statistical testing step. This figure is an adjusted version of Figure 1 in Mieth et al.^31^

The main motivation behind DeepCOMBI is to harness the immense potential of sophisticated, state-of-the-art artificial intelligence (AI) methods to examine complex and potentially nonlinear structures in high-dimensional data by applying the concept of DNNs to GWAS in the first step of the algorithm. Subsequently, in step 2, DeepCOMBI identifies a set of SNPs that have high effects on the classification result of the DNN (either individually or in combination with other SNPs and not due to correlation structures) by calculating an explanation score for each SNP which reflects its contribution to the final classification decision. The third and last step assigns individual *p*-values to all selected SNPs and quantifies its relevance with a permutation based significance threshold.

**Figure 1** gives an overview of the overall workflow of the DeepCOMBI method, which is described in detail in the **Methods Section**. DeepCOMBIs performance on both controlled generated datasets as well as on a 2007 GWAS dataset of seven common diseases^14^ is presented in the **Results Section.** We show that DeepCOMBI compares favorably to a number of competitor methods in terms of both classification accuracy as well as SNP association prediction when validated with all associations reported within the GWAS catalog accessed in 2020. A thorough discussion of the results and all related machine learning work is given in the **Discussion Section.** An implementation of the DeepCOMBI method in Python is available on github at https://github.com/AlexandreRozier/DeepCombi.

## Methods

The proposed method applies deep learning and the concept of explainable AI to GWAS data and enables the identification of SNPs that are associated with a given trait with statistical significance. A graphical representation of the method is given in **Figure 1**. The method is based on a deep learning step that trains a DNN for the classification of GWAS subjects into their respective phenotype class. Using LRP as a post-hoc explanation method, we access the relevances of all SNPs regarding each individual classification result. The obtained SNP relevance scores are used to select the subset of most important SNPs to test for association in the final multiple testing step.

In the following sections we describe the statistical problem, which is investigated in a GWAS, present the proposed method in detail and specify the experimental setup of performance assessments on generated synthetic data and a real-world application of a known GWAS dataset.

### Problem setting

A GWAS investigates the observed genotypes 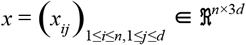 of *d* SNPs and *n* subjects labelled with the corresponding phenotypes *y* = (*y*_1_..., *y_n_*). Both the genotypic information in SNP *j* of subject *i* and the phenotypes are encoded in a binary way, where *x*, ∈ {(1,0,0),(0,1,0),(0,0,1)} represents the number of minor alleles and *y*, ∈ {0,1} is the binary label separating controls from cases. The null hypothesis of a conventional single locus test is that there is no difference between the trait means of any genotype group, which would indicate that the genotype at SNP *j* is independent of the phenotype under investigation^54^. Via a chi-square test RPVT calculates a *p*-value *j* for each SNP *j* and declares it significantly associated with the phenotype if *pj,* ≤ *t\**. The threshold *t\** has to be chosen carefully as the significance level α in the case of a single test and adjusted if multiple tests are being conducted to bound the family-wise error rate (FWER), i.e. the probability of at least one false positive test result, to α. Bonferroni correction is the most straightforward way to take multiplicity into account by setting 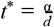. ^61^

The individual RPVT *p*-value for association of the *j*-th SNP only depends on *x_*j_* and thus disregards any possible correlations and interactions with other SNPs. Additional information can be yielded by applying machine learning based prediction methods which use the information of the whole genotype and calculating *p*-values only for the SNPs that were of importance in the decision process of such machines.

### DeepCOMBI

Combining the concepts of DNNs, explanation methods and statistical testing, we propose a novel algorithm consisting of the following three steps:

#### 1. Deep learning

Given the genotypes *x* = (*x_ij_*) and the corresponding phenotypes *y* = (*y_i_*,) of a GWAS - a DNN is trained for phenotype prediction.

#### 2. Explanation and SNP selection

A subset of SNPs is selected by applying LRP as an explanation method for each individual prediction and averaging the absolute values of the resulting explanations to compute global prediction relevance scores *r*_1_,…, *r_d_*. The relevance scores are processed through a moving average filter with window size *l* and given a predefined upper bound *k*∈ {1,…,*d*} for the number of informative SNPs, we select the *k* most relevant SNPs based on *r.*

#### 3. Statistical testing

A hypothesis test is performed for all SNPs selected in the previous step to compute *p*-values of those SNPs, while the *p*-values of all other SNPs are set to one. Via a permutation based threshold calibration and given a FWER level *α*, we decide that SNP *j* is associated with the trait if *p_j_* < *t\**, where *t*\* ≡ *t*\* (*k*, α) is chosen as the α-quantile of the permutation distribution of the *k* smallest *p*-values.

The proposed algorithm can be viewed as an extension of the COMBI method^31^, a two-step method including an SVM step and a statistical testing step. We replace the former with state-of-the-art deep learning methods and explanation techniques.

The above steps are presented in detail in the following sections.

##### The first step of DeepCOMBI - Deep learning

The first step of the proposed method consists of constructing and training a well-performing DNN for the prediction of the phenotypes *y* = (*y_i_*)of a GWAS given the corresponding genotypes *x* = (*x*). Selecting a DNN architecture is often critical for achieving good performance for a specific - in this case SNP-based - classification task. Montaez et al.^43^ developed a 2-class DNN for the classification of polygenic obesity and have successfully shown its performance to be superior to numerous competitor methods. Romagnoni et al.^28^ have compared the performance of similar architectures and have presented a detailed review of the best design choices for a DNN on a Crohn’s Disease dataset. Taking inspiration from the conclusions of both of these works and having checked performances on synthetic datasets, we use an architecture of two fully connected layers with 64 neurons and ReLU activations and a dense softmax output layer with two output nodes. To improve validation accuracy by reducing overfitting, each hidden layer is followed by a dropout layer with a dropout probability of ϕ.

The loss function to be optimized in the training process is based on the classic cross entropy loss. To guarantee good generalization to unseen samples and avoid overfitting despite the large number of model parameters, the binary cross entropy loss is coupled with an L1-L2 mixed regularization term:

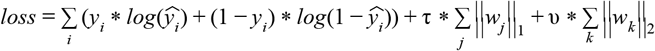

with *y_i_*. being the ground truth label, 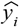 the predicted class which depends on the learned parameters *w* of the DNN and *τ,ν*>0 the regularization parameters. This loss function guarantees that the network avoids overfitting by minimizing the trade-off between small errors on the data on the one hand and small L1 and L2 norms of the vector *w* on the other hand. Adam^55^ is used as an adaptive learning rate optimizer to minimize the given loss function.

To overcome limitations due to imbalanced datasets, class weights were calculated according to the class frequencies and used to direct the DNN to balance the impact of controls and cases.

Once the parameters *w* of the DNN have been trained by optimizing the above learning problem the network is able to predict the phenotype of any unseen genotype *x*. Regarding this binary classification problem, the output node with the highest score represents the predicted phenotype.

In a preprocessing step the data is centered and scaled by subtracting the global mean and dividing by the global standard deviation. To minimize computational effort and limit the number of model parameters in the DNN a *p*-value threshold κ can be applied in order to only select SNPs with *p*-values smaller than κ to be used for training.

##### The second step of DeepCOMBI - Explanation and SNP selection

To harness the potential of DNNs in the identification of SNP disease associations in GWAS we now apply the concept of explainable AI. Once the DNN is fully trained, the aim is to define an importance measure that determines which loci play an important role in the determination of a phenotype. Generating relevance scores from trained DNNs can be achieved by using LRP^48,49,50^, which consists of the following two steps: After a DNN *f* is trained on a prediction task, the prediction scores of a datapoint *x_i_* are computed by *f*(*x_i_*) = *y_i_*, a forward pass through the network. Afterwards, following a specific propagation rule, a single output score, i.e. the highest output score, *y_i_* is backpropagated successively layer-by-layer through the network until reaching the input layer. In this work, we use the αβ - LRP rule, where the relevance 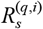 of neuron *s* in layer *q* depends on the relevance of all of its successors *t* in layer *q+1* in the following way:

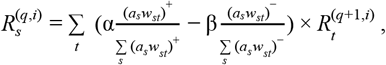

where *a_s_* denotes the activation of neuron *s*, and *w_st_* is the weight between the two neurons *s* and *t*. This rule allows us to weigh the positive and negative contributions of neurons *t* to their predecessor *s* differently by α and β.

Once the input layer 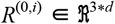 is reached, a relevance score ρ_*ij*_ of SNP *j* in subject *i* is attributed to each dimension of *x_i_* with 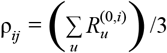. Since the original relevance vector *R*^(0, *j*)^ contains three values for each one hot encoded location, it is converted back to size *d* by averaging over the three nodes *u* ∈ {(*j* × 3)-2, (*j* × 3)-1, (*j* * 3)} corresponding to SNP j in the input layer.

Note, that all relevance scores ρ_*ji*_ will be positive, since a softmax output layer with two output nodes for the binary classification problem was used and only the highest of the two output activations was backpropagated.

ρ_*ij*_ now demonstrates to which extent the dimension *j* of *x_i_* plays a role in the classification decision *f*(*x_i_*) and can be used to uncover the most relevant SNPs for prediction. Note however, that LRP is applied individually to each datapoint *i*. By averaging the values of all individual LRP explanations ρ_*ij*_ of SNP *j* we propose to generate a global explanation 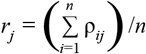, which is independent of data points. The relevance scores of one sample sum up to the activation value of the output prediction, which means that data points classified with low certainty will also have a small impact on the global explanation. Intuitively, the global LRP score *r_j_* of each SNP *j* can now be interpreted as a measure of relevance regarding the prediction. The higher *r_j_*, the greater the influence of locus *j* on the decision process of the DNN.

To achieve better performance, Mieth et al.^31^ suggested that SNP relevance scores should be filtered before using them to select the highest scoring locations. Hence, the LRP relevance score vector *r* is post-processed through a *p*-th-order moving average filter, that is:

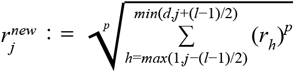

where *l*∈1,…,*d* denotes the window size *l* and *p*∈]0,∞[. We have now generated relevance scores showing which SNPs played an important role in the classification decision and can use them for the selection of promising locations. For the next step of DeepCOMBI we choose to test all SNPs with the *k* largest values of the scores 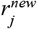 and eliminate all SNPs with lower relevance.

##### The third step of DeepCOMBI - Statistical testing

The Statistical testing step of the DeepCOMBI method is directly derived from the second step of the COMBI method^31^. A χ^2^ hypothesis test is performed for each of the *k* SNPs selected in the LRP explanation step and the *p*-values for all other SNPs are set to one. To identify statistically significant associations, a *p*-value threshold *t*\* is calibrated to control the *FWER* for multiplicity by applying the permutation procedure proposed by Mieth et al.^31^. They developed an extension of the Westfall and Young procedure^56^. A thorough discussion and derivation of the method, its assumptions and validity generally and in this specific application can be found here^56,57,58^ and here^31^, respectively. We estimate the distribution of *p*-values under the global null hypothesis of no informative SNPs by repeatedly assigning a random permutation of the phenotypes to the observed genotypes and applying the complete workflow of the DeepCOMBI method to save the resulting *p*-values of the *B* Monte Carlo repetitions^31^. “The empirical lower α -quantile of the smallest of these *p*-values is then a valid choice for *t\** in the sense that the *FWER* for the entire procedure is bounded by α ”^31^. In contrast to the Bonferroni threshold calibration this procedure takes all dependencies in GWAS datasets caused by strong linkage disequilibrium (LD) into account.

### Baselines

In order to evaluate the performance of the proposed DeepCOMBI method in comparison to competitor approaches, we select a set of representative baseline methods. RPVT is chosen as the most widely used traditional, purely statistical testing approach. As a machine learning based method and the methodological background of DeepCOMBI we select COMBI as the main competitor method we aim to succeed in terms of performance. Since the COMBI method was shown to outperform other combinatorial machine learning based approaches (Roshan et al.^6^, Meinshausen et al.^58^ and Wasserman and Roeder^59^) and a number of purely statistical analysis tools (Lippert et al.^21,27^) on the same datasets evaluation methods used here, there is no need to compare to those methods again.

#### RPVT as baseline

Raw *p*-value thresholding (RPVT) is a statistical framework traditionally used in GWAS for identifying significant associations between SNPs and traits. The single locus null hypothesis to be tested states that the SNP at locus *j* is independent of the binary trait of interest, i.e. that there is no correlation between this particular SNP and the development of the disease under investigation. A standard statistical test for this hypothesis is the χ^2^-test^60^, which tests for independence of the two multi-level variables genotype (three different levels: 0, 1 or 2 minor alleles) and phenotype (two different levels: case or control) by calculating the test statistic 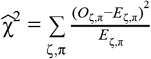 where *O_ζ,π_* and *E_ζ,π_* are the observed and expected frequencies of genotype *ζ* in combination with phenotype *π*. To compute a *p*-value 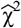 is then compared to a χ^2^ distribution with two degrees of freedom and represents the probability of observing a sample statistic as extreme as 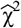 under the assumption of no association between genotype and phenotype. If the *p*-value is smaller than a predefined threshold *t*\* the null hypothesis is rejected and we declare the SNP under investigation to be significantly associated with the phenotype. If there was a single test to perform *t*\* would usually be equal to the significance level α = 0.05. When performing multiple testing, however, the threshold is modified to take the multiplicity of the problem into account. The simplest method is the so-called Bonferroni correction^61^, where *t\** is divided by the number of tests performed, i.e. *d*, the number of SNPs in our case, which guarantees that the FWER, the probability of one or more erroneously reported associations, is bounded by α. The Bonferroni correction works well under the assumption that all null hypotheses are independent of each other, which is not the case here. Indeed, since SNPs show high degrees of correlation through LD, the Bonferroni correction can become extremely conservative leading to a high rate of false rejections, which is why the scientific community mostly applies a fixed threshold that remains constant for multiple GWAS. Here, based on the original publication of the data we are analyzing (WTCCC data, see the **Methods Section** on validation datasets^14^) and the findings of Mieth et al.^31^, we present not only the strong associations at a significance level of *t\** = 5*x*10^−7^ but also weak associations at *t\** = 1 × 10^−5^.

#### COMBI as baseline

The COMBI method^31^ combines machine learning with multiple hypothesis testing to improve the statistical power of GWAS. It is a two-step method including the training of a SVM^53^ and using the resulting SVM weights as importance scores to select a subset of candidate SNPs for statistical testing. In the first step of COMBI an SVM for the prediction of the unknown phenotype *y* based on the observation of genotype *x* is trained to determine the weight vector *w.* The following optimization problem is solved:

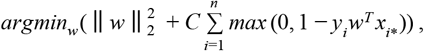

where *C* is the regularization parameter that controls the trade-off between a small norm of *w* and a small prediction error of the machine. After training, the weight vector *w* is filtered and interpreted as an importance score to determine which loci play an important role in the decision process of the SVM. A χ^2^ test is performed only on the SNPs with the highest scores while all other *p*-values are set to one. The same permutation test procedure as described in the **Methods Section** about the multiple testing procedure of DeepCOMBI is applied to define a significance threshold *t\**.

#### Raw SVM weights and LRP scores without statistical testing as baselines

Instead of interpreting the SVM weights from COMBI and the LRP scores from DeepCOMBI as relevance scores to select a subset of SNPs to calculate *p*-values for, this step can be skipped to use the raw SVM and LRP scores as a test statistic. For evaluation the vector of raw SVM weights and LRP scores can be treated like the vector of *p*-values of RPVT, COMBI and DeepCOMBI to calculate performance curves. We compare DeepCOMBI to these baseline methods of raw relevance scores and RPVT to show that only the combination of machine learning / deep learning and multiple testing show the desired performance increase which cannot be achieved individually by one of the components.

### Validation datasets

#### Validation on generated datasets

To create a realistic but controlled environment where the ground truth labels of a dataset, i.e. the SNPs that are indeed linked to the disease, are known, we generate semi-synthetic data for a first evaluation of DeepCOMBI and the baseline methods from above. We follow the instructions for the creation of such GWAS datasets proposed by Mieth et al.^31^ The basic concept is to take an ensemble of real genotypes and generate a synthetic phenotype for each subject according to a specific rule. With this method, the underlying architecture of the genome, including for example genetic LD and correlation structures, is kept intact while control over the phenotypic labels is gained at the same time.

We use the WTCCC dataset^14^ described in more detail below and randomly select 300 subjects of the Crohn’s disease dataset. We draw a random block of 20 consecutive SNPs from chromosome 1 and a random block of 10,000 consecutive SNPs from chromosome 2. The former are representing the informative SNPs and are placed in the middle of the 10,000 uninformative SNPs. Synthetic phenotypes are now generated only based on one of the informative SNPs (at position 5010) according to the following phenotype probability distribution:
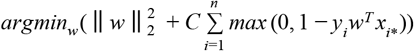
where *γ* is an effect size parameter, *x_i,*_* is the allele sequence in nominal feature encoding (i.e. *x_ij_* is the number of minor alleles in SNP *j* of subject *i*) and *Y_i_* is the generated phenotype of subject *i*. Basing the label of a subject on the SNP at position 5010 alone, will create associations to all 20 informative SNPs and typical tower shaped *p*-value formations in the resulting Manhattan plots, because there are real covariance structures and LD within the 20 informative SNPs. At the same time the tower structure is limited to those 20 informative positions, because there are no correlations of those 20 SNPs with the surrounding 10,000 noise SNPs coming from chromosome 2. The random generation process will also ensure that the datasets will have associations of different strengths to the 20 informative SNPs. The complete data generation process is repeated to generate 1,000 datasets. DeepCOMBI and all baseline methods are applied to each dataset with an 80:20 class balanced split in training and test data. The prediction results on the test data are evaluated with the known ground truth of only 20 informative SNPs at the positions 5000 to 5020 and the corresponding performance can be measured in terms of the number of true and false positives for each method.

#### Validation on WTCCC data

For evaluation on real-world genomic data the performance of DeepCOMBI was assessed on the Wellcome Trust Case Control Consortium phase 1 dataset, released in 2007^14^ featuring the genotypic information on 500,000 SNPs of 17,000 British subjects. With 3,000 shared controls and 2,000 case samples for seven major human diseases (Crohn’s disease (CD), type 1 diabetes (T1D), type 2 diabetes (T2D), coronary artery disease (CAD), hypertension (HT), bipolar disorder (BD) and rheumatoid arthritis (RA)) it was a landmark study both in terms of sample size and dimensionality at the time of its publication. For our analysis a case-control dataset for each disease was created removing all SNPs and samples that did not fulfill the quality control criteria provided in the original WTCCC paper.

In agreement with the lack of inter-chromosomal LD and Mieth et al.^31^ who showed no significant performance increase with genome wide training, the DeepCOMBI method and all baseline methods were applied to each chromosome separately.

For evaluation purposes the concept of replicability was applied based on Mieth et al.^31^. Since the true underlying genetic architecture of the given traits, i.e. the sets of informative SNPs for each disease are unknown, an approximation of the truth was created by employing the GWAS catalog^62^ and examining the results of the 13 years of independent studies after the WTCCC dataset was published. To evaluate the reported finding of a method (DeepCOMBI or competitor) the GWAS catalog (accessed on July 30, 2020) is inquired for that SNP and all SNPs in LD ((R^2^>0.2) according to PLINK LD calculations^63^) within a 200kb window around that SNP. If an association with the disease with *p*-value <10^−5^ of the SNP itself or the SNPs in LD was reported by at least one independent GWAS published after the WTCCC study, the reported SNP is counted as a true positive finding. In contrast, all SNPs that were not replicated count as false negatives.

### Parameter selection

The application of DeepCOMBI requires the determination of a number of free parameters. In the following sections we present the selected optimal parameter values and describe the process of finding them for the different datasets under investigation.

#### Parameter selection on generated datasets

For the generation process of semi-synthetic datasets all parameters were selected according to the information given in Mieth et al.^31^. Most importantly, the effect size parameter was set to γ = 6.

When applying the DeepCOMBI method to the generated datasets, we studied the effect of all hyperparameters on the performance of the DNN. An accuracy based random grid search with a stratified split in 90% training and 10% testing data was conducted. Here, we present the selected most successful values and the investigated parameter intervals in parentheses:

- number of neurons per dense hidden layer nn = 64 [2, 4, 8, 16, 64],
- L1 regularization coefficient τ = 0.0001 [0, 1e-6, 1e-5, 1e-4, 1e-3,1e-2, 1e-1],
- L2 regularization coefficient ν = 0.000001 [0, 1e-6, 1e-5, 1e-4, 1e-3,1e-2, 1e-1],
- dropout rate ϕ =0.3 [0.3, 0.5],
- learning rate η = 0.01 with learning rate reduction on plateau with factor 0.7125 after 50 epochs of no improvement,
- number of epochs e = 500 [100, 500, 1000].

A few different parameter values of the αβ - backpropagation rule were manually investigated on exemplary datasets. By visually inspecting the resulting LRP vectors and their corresponding DeepCOMBI *p*-values the combination of α = 1 [0, 1, 2] and β = 0 [0, 1, 2] was found to be best.

For post processing the global relevance scores and selecting the most relevant SNPs, we assumed that the most successful values found by Mieth et al.^31^ would also be a good choice for our method. Hence, we set the window size of the moving average filter to *l* = 35, the norm parameter of the moving average filter to */* = 2 and the SNP selection parameter to *k* = 30. These values were found to be in agreement with the biological background of the data, e.g. *l* = 35 reflects the reach of LD along a genetic sequence.^31^

#### Parameter selection on WTCCC data

To choose hyperparameters for the DNN trained on WTCCC data in the first step of DeepCOMBI, a parameter search was run on a single dataset. The Crohn’s disease chromosome 3 dataset was selected as a good representative and an accuracy based parameter search with a stratified split in 90% training and 10% testing data was conducted. We studied the effect of the hyperparameters on the performance of the DNN and the best performing hyperparameters were as follows (tested intervals in parentheses):

- number of neurons per dense hidden layer *nn* = 64 [2, 4, 8, 16, 64],
- L1 regularization coefficient τ = 0.001 [0, 1e-6, 1e-5, 1e-4, 1e-3,1e-2, 1e-1],
- L2 regularization coefficient ν = 0.0001 [0, 1e-6, 1e-5, 1e-4, 1e-3,1e-2, 1e-1],
- dropout rate ϕ = 0.3 [0.3, 0.5],
- *p*-value threshold κ= 1e-2 [1e-4, 1e-2, 1],
- learning rate η = 0.00001 [1e-7, 1e-6, 1e-5, 1e-4, 1e-3,1e-2, 1e-1],
- number of epochs *e* = 500 [100, 500, 1000].

Detailed results on the classification performance of the final training parameter settings can be found in the **Results Section**.

As before, we visually investigated a few different parameter values of the αβ - backpropagation rule and their influence on both the resulting relevance scores and *p*-values. On the Crohn’s disease chromosome 3 dataset the combination of α = 2 [0, 1, 2] and β = 1 [0, 1, 2] was found to be optimal.

After manually investigating the global LRP scores and the corresponding DeepCOMBI */-*values of the exemplary dataset (Crohn’s disease chromosome 3), we found that slightly different settings than for the analysis of the generated datasets should be applied for post processing the relevance vectors and selecting the most relevant SNPs. Namely, the window size of the moving average filter should be set to *l* = 21 and the SNP selection parameter should be increased to *k* = 200. The need for a decreased filter size and an increased number of selected SNPs might be caused by the application of the *p*-value based pre-selection step for limiting the number of model parameters, which is only applied to the real dataset and not the generated datasets.

To determine the value of the significance level α to be used in the permutation test procedure of the last steps DeepCOMBI, we follow the recommendations of Mieth et al.^31^ who calculated the empirical distribution of *p*-values using the Westfall-Young^56^ procedure and determined the error level that the RPVT threshold of *t\** = 1 × 10^−5^ corresponds to. For a valid comparison to both the original WTCCC study as well as the COMBI publication we employ the same significance levels.

All free parameters of the COMBI method, e.g. the SVM optimization parameter *C,* were set according to the original COMBI publication^31^.

### Performance metrics

To assess the performance of DeepCOMBI and the baseline methods a number of statistical metrics were evaluated. The performances of both the intermediate step of classification (of SVMs and DNNs) and the final result of predicted informative SNPs need to be explored.

Assuming we know the ground truth, the metrics are defined as follows:

- TP = True positive; FP = False positive; TN = True negative; FN = False negative
- Accuracy = (TP + TN)/(TP + TN + FP + FN)
- Balanced accuracy = (TPR + TNR)/2
- Precision = TP/(TP + FP)
- True positive rate TPR = TP / (TP + FN)
- False positive rate FPR = FP / (TP + FN)
- Family-wise error rate FWER = P(FP >=1)

The following performance curves and the area under these curves (AUC) will be investigated:

- Receiver operating characteristic curve (ROC): TPR vs. FPR or TP vs. FP or TPR vs. FWER
- Precision-recall curve (PR): Precision vs. TPR or Precision vs. TP

### Implementation details

The DeepCOMBI method was implemented in Python and the source code is available at https://github.com/AlexandreRozier/DeepCombi. The implementation uses the DNN development library Keras^64^ in combination with the LRP library iNNvestigate^65^.

## Results

In the following section we present the results of the proposed DeepCOMBI method evaluated on generated as well as on real world data. Performance in terms of both classification accuracy and SNP prediction is examined in comparison to a number of baseline methods, which are presented in full detail in the **Methods Section**. As evaluation criteria, we report prediction accuracy for the classification step and *FWER,* precision and *TPR* for the SNP selection step. See the **Methods Section** above for a detailed description of the evaluated performance metrics.

### Results on generated datasets

Here, we report our results averaged over the 1,000 data sets generated in the simulation process described in the **Methods Section** (“Validation on generated datasets”). We show that on these data sets DeepCOMBI performs better than the traditionally used method for analyzing GWAS, RPVT, and its main competitor, the COMBI method.

#### Prediction performance on generated datasets

The first steps of both DeepCOMBI and COMBI consist of training a learning algorithm for the classification of all subjects into their respective phenotypic group given their genotypic information. Since all following steps depend on the performance of these classifiers, high prediction accuracy is crucial. On the generated datasets the SVM (as part of the COMBI method) achieves 59% accuracy on average and 54% balanced accuracy. In comparison, the DNN (as part of the DeepCOMBI method) performs significantly better with an average of 74% classification accuracy and also avoids negative effects of unbalanced datasets more effectively by applying class weights in the DNN training (74% balanced accuracy). Accuracy scores and additional information are given in **Table 1**. Following these promising intermediate results, in the next section we investigate whether the entire workflow of the DeepCOMBI method can also outperform the baselines methods.

**Table 1.** Classification performance on generated datasets. Summary statistics of the classification accuracy of the SVM (as in the first step of COMBI) and the DNN (as in the first step of DeepCOMBI) are presented. Values corresponding to accuracy and balanced accuracy in parentheses are given.

|  | Mean accuracy<br>(balanced accuracy) | Standard deviation of accuracy<br>(balanced accuracy) | Minimum of accuracy<br>(balanced accuracy) | Maximum of accuracy<br>(balanced accuracy) |
| --- | --- | --- | --- | --- |
| **SVM (as in COMBI)** | 0.59 (0.54) | 0.05 (0.06) | 0.41 (0.35) | 0.76 (0.71) |
| **DNN (as in DeepCOMBI)** | 0.74 (0.74) | 0.07 (0.07) | 0.55 (0.50) | 0.97 (0.98) |

#### SNP selection performance on generated datasets

To compare the relevance scores and *p*-values obtained with the novel LRP-based method to those derived from the SVM weights in the COMBI method we look at three exemplary synthetic datasets and the corresponding results (See **Figure 2**). They can be distinguished by the level of association of the 20 informative SNPs (highlighted in pink) with the phenotype. In the first column of subfigures the strength of association for each replication at positions 5001 - 5020 is shown in the corresponding RPVT Manhattan plots. While the first row of subfigures represents a replication with very weak associations (small pink tower), the second has a moderate association (medium sized tower) and the third shows a very strong association (large tower). In the second and third column the raw SVM weights and LRP scores are shown. It can be seen that LRP yields clearer relevance distributions in comparison to the SVM-based method. Even with the huge number of parameters the DeepCOMBI models explanation score yields a lot less noise than the SVM weights of COMBI. This results in the COMBI method only being able to classify the very strong association correctly (third column of subfigures), while it misses the weak and moderate ones. In contrast, DeepCOMBI is successful for both the second and third replication with moderate and strong associations and only misses the very weak association (last column of subfigures). Please note that DeepCOMBI not only precisely identifies the correct informative tower, but also filters out a relatively high noise tower at around position 600 which - just by chance - achieved a *p*-value < 10^−5^. The method thus not only increases the probability of finding the correct tower but also, and potentially more importantly, decreases the probability of falsely selecting a noise tower^31^.

**Figure 2:**
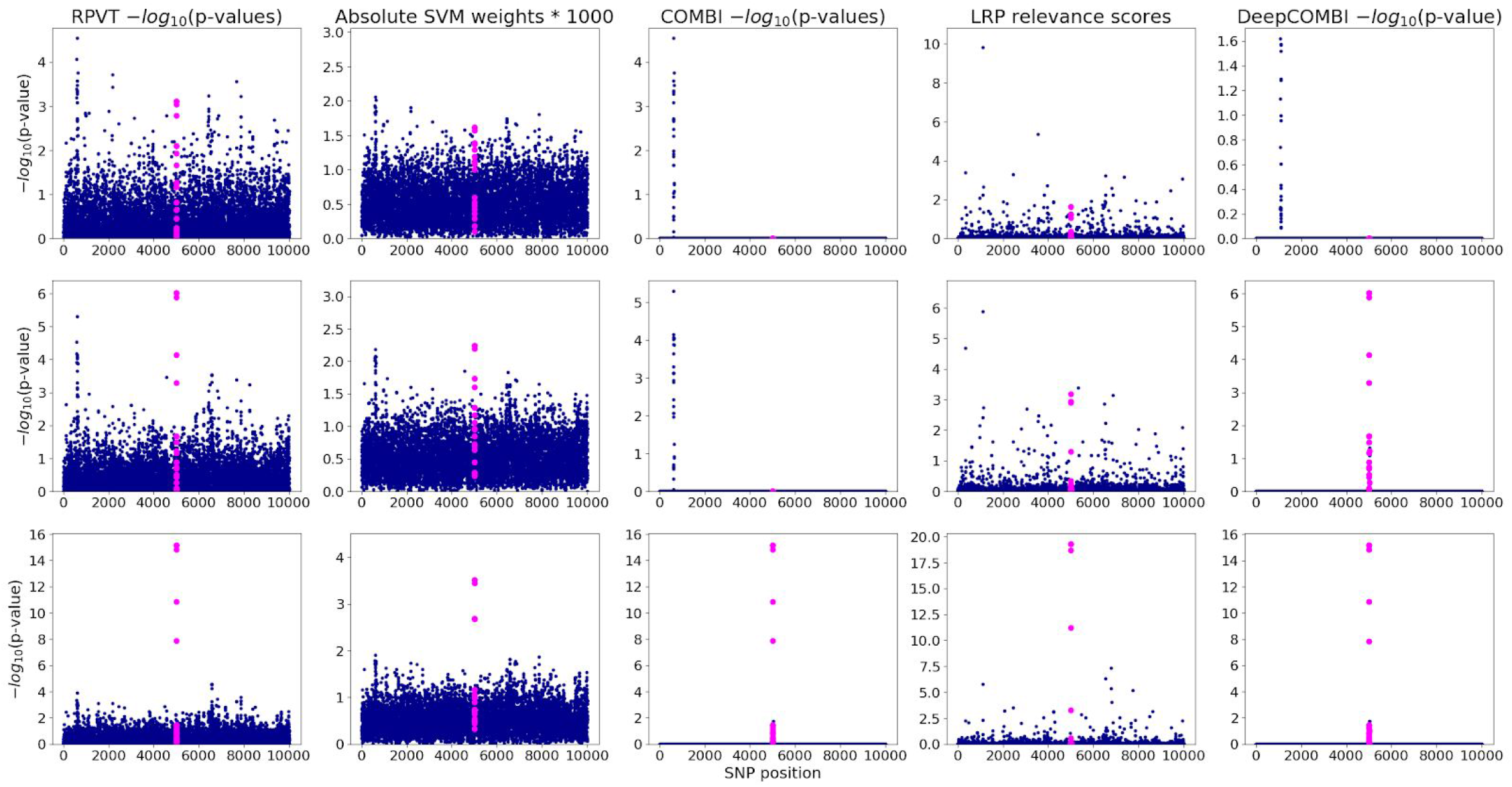
Three exemplary generated datasets and the corresponding COMBI and DeepCOMBI results. We present the results of three exemplary replications: one with weak (first row), one with medium (second row) and one with strong (third row) association of the 20 informative SNPs at position 5001-5020 (highlighted in pink in all subfigures). Standard RPVT *p-*values are plotted in the first column of subfigures. Absolute SVM weights and corresponding *p-*values of the COMBI method are shown in the second and third column. Finally, LRP relevance scores and the corresponding *p-*values of DeepCOMBI are presented in the fourth and last column.

To investigate whether these exemplary findings represent a general trend, we now examine the results of all competitor methods averaged over all 1,000 generated datasets. In **Figure 3** the corresponding ROC and PR curves are shown. In both subfigures and for all levels of error and detection rates DeepCOMBI (pink line) consistently outperforms RPVT (light blue line) and COMBI (dark blue line) in terms of power and precision. The combinatorial approaches, DeepCOMBI and COMBI, also perform better than their individual components of a machine learning algorithm (SVM or DNN with LRP) and a multiple testing step (RPVT). This can be deduced from the fact that RPVT as well as the other two baseline methods of directly thresholding the raw LRP scores (dashed pink line) and SVM weights (dashed dark blue line) separately cannot achieve the same performance as their combinations (i.e. DeepCOMBI and COMBI).

**Figure 3:**
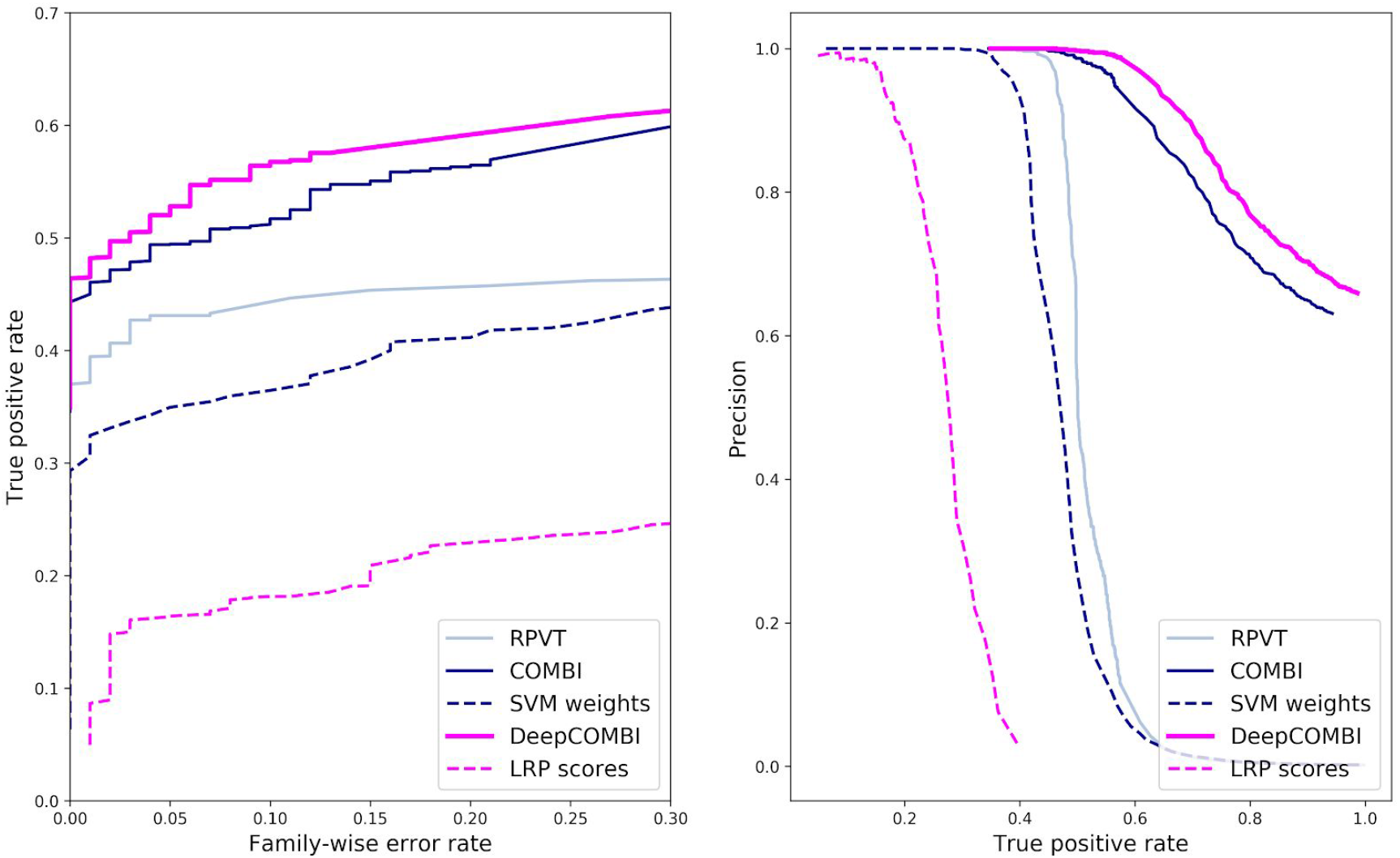
ROC and PR curves of DeepCOMBI and all competitor methods on generated datasets. Performance curves of all methods averaged over the 1,000 generated datasets are shown. ROC curves are presented on the left and PR curves on the right side.

### Results on WTCCC data

#### Prediction performance on WTCCC data

In the first step of DeepCOMBI a DNN is trained and we investigate its performance on all diseases and chromosomes. **Figure 4** shows that the DNN of DeepCOMBI performs consistently better than the SVM of COMBI in terms of all four validation metrics described in the **Methods Section** under “Performance metrics”.

**Figure 4.**
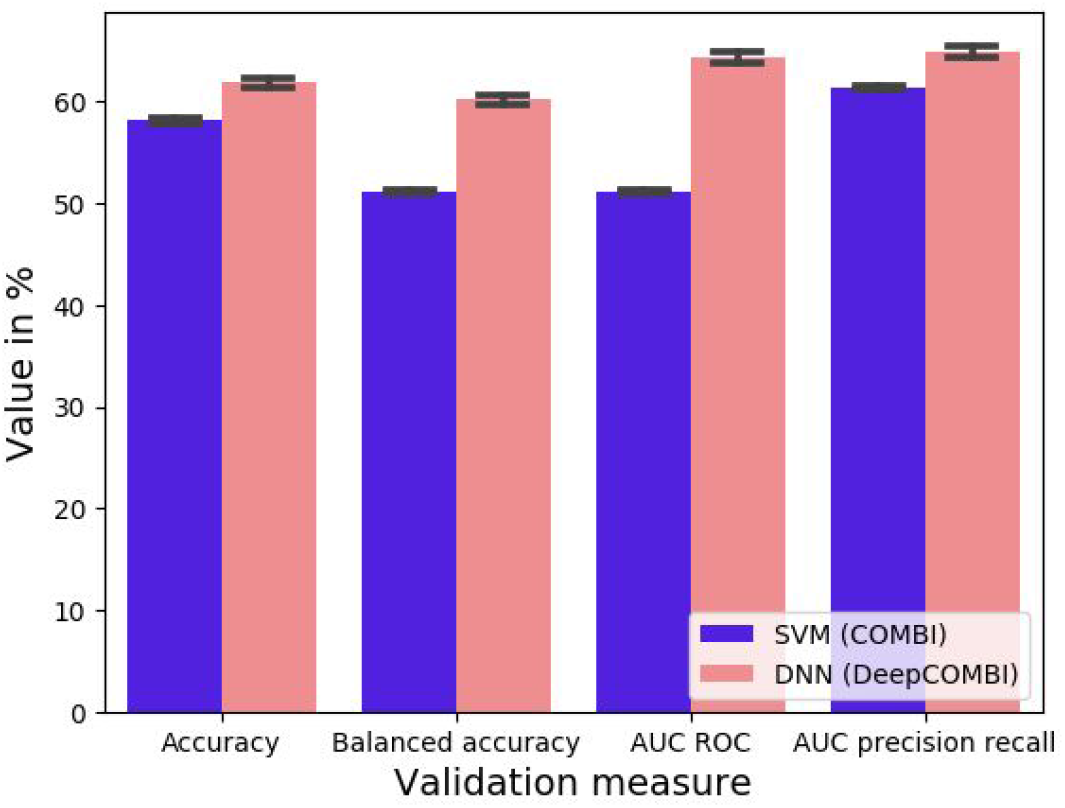
Classification performance on WTCCC data. Mean validation measures of SVM (as in the first step of COMBI) and DNN (as in the first step of DeepCOMBI) averaged over all diseases and chromosomes are given with standard deviation. All datasets were split into 80% training and 20% validation data.

##### SNP selection performance on WTCCC data

In **Figure 5** we present the results of the traditional RPVT approach, the COMBI method and the DeepCOMBI applied to the seven diseases of the WTCCC 2007 dataset. In each corresponding Manhattan plot the negative logarithmic *p*-values of all SNPs at a given position in a chromosome are shown. While RPVT assigns *p*-values smaller than one (i.e. nonzero in the plots on logarithmic scale) to all SNPs and in consequence produces a lot of statistical noise, both COMBI and DeepCOMBI discard most SNPs (i.e. assign *p*-value one, i.e. zero in the plot on logarithmic scale) and hence reduce the level of noise significantly. The COMBI method selects 100 SNPs with high SVM weights per chromosome and DeepCOMBI chooses 200 SNPs with high LRP scores. In all plots the significance threshold *t*^*^ is represented by dashed horizontal lines and all statistically significant SNP associations are highlighted in green. Please note, that in the case of RPVT the threshold is constant at *t\** = 1 × 10^−5^ (i.e. 5 in the plot) for all chromosomes. A chromosome wise threshold was generated for both COMBI and DeepCOMBI via the permutation based procedure described in the **Methods Section** to match the expected number of false rejections of RPVT.

**Figure 5.**
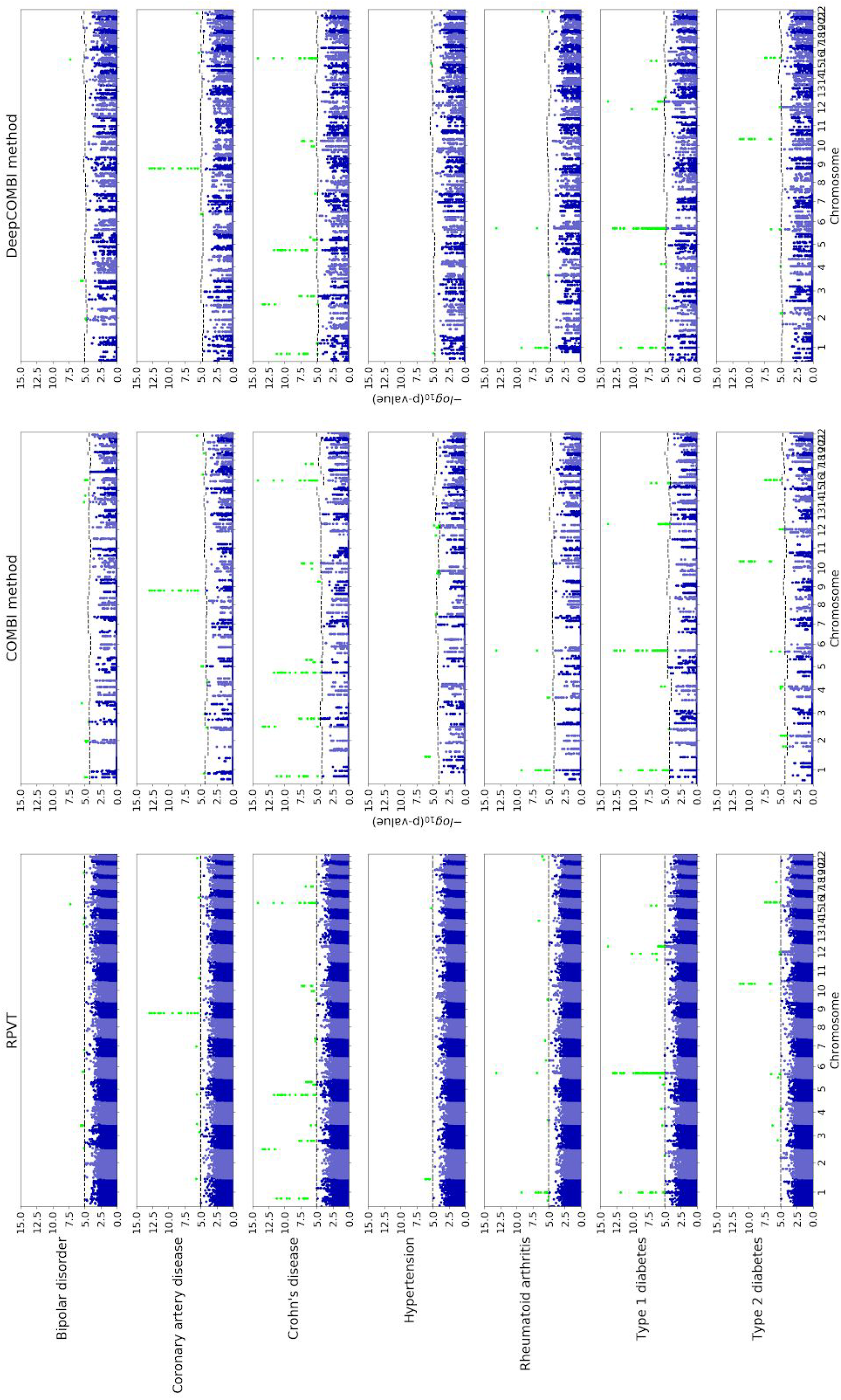
Manhattan plots for WTCCC data. The negative logarithmic χ^2^ test *p*-values are plotted against position on each chromosome for all seven diseases. Results from the standard RPVT approach, the COMBI method and the DeepCOMBI method are shown. Thresholds indicating statistical significance are represented by dashed horizontal lines and significant *p*-values are highlighted in green. Please note that the y-axes of all plots have the same limits (0 to 15) to enable direct comparison.

All SNPs reaching statistical significance in the permutation-based thresholding procedure of the DeepCOMBI method are presented in **Table 2**. Besides showing basic information (associated disease, chromosome, identifier and χ^2^ *p*-value) for all of these SNPs, the fifth and sixth column indicate whether they were found to be significant by RPVT with the application of *t\** = 10^−5^ or by the COMBI method. To validate all findings, the seventh and eighth column report whether - and if so in which external study - they have been found significantly associated with the given disease according to the GWAS catalog. By investigating whether the identified SNPs were discovered as significant in an independent GWAS published after the original WTCCC study it can be determined to which extent those novel findings can be confirmed to be true associations.

**Table 2:** Significant SNPs of the DeepCOMBI method and related association details. For each SNP identifier on a specific chromosome that was found to be significantly associated with a disease by the DeepCOMBI method, we show their χ^2^ test *p*-value and indicate whether the RPVT *p*-value is < 10^−5^ (i.e. the SNP is a significant finding of RPVT), whether its COMBI *p*-value is smaller than the corresponding COMBI threshold (i.e. the SNP is a significant finding of the COMBI method) and whether the SNP has been found significant with a *p*-value < 10^−5^ in an external study with a corresponding PMID. Please note that the RPVT result in the fifth column corresponds to the χ^2^ *p*-values we have calculated here, not necessarily to the original WTCCC publication, where they also investigated trend test *p*-values and potentially applied slightly different pre-processing steps. Similarly, the COMBI result in the sixth column corresponds to the re-calculations of COMBI we performed here, not necessarily to those of the original COMBI publication where slightly different results were produced due to the random nature of the permutation procedure.

| Disease | Chromosome | Identifier | $\chi^2$ $p$-value | Significant in RPVT | Significant in COMBI | $p$-value $< 10^{-5}$ in at least one external GWAS or meta-analysis | References (PMID) |
| --- | --- | --- | --- | --- | --- | --- | --- |
| Bipolar disorder (BD) | 2 | rs7570682 | 1.77e-05 |  | YES | YES | 21254220 |
|  | 2 | rs1375144 | 1.26e-05 |  | YES | YES | 21254220 |
|  | 3 | rs514636 | 2.53e-06 | YES | YES | YES | 21254220 |
|  | 16 | rs420259 | 5.87e-08 | YES | YES | YES | 21254220 |
| Coronary artery disease (CAD) | 6 | rs6907487 | 2.92e-05 |  |  | YES | 17634449 |
|  | 9 | rs1333049 | 1.12e-13 | YES | YES | YES | 17634449 |
|  | 16 | rs8055236 | 5.32e-06 | YES | YES |  |  |
|  | 22 | rs688034 | 2.75e-06 | YES | YES |  |  |
| Crohn's disease (CD) | 1 | rs11805303 | 6.35e-12 | YES | YES | YES | 17435756 |
|  | 1 | rs12037606 | 1.02e-05 |  |  | YES | 17554261 |
|  | 2 | rs10210302 | 4.52e-14 | YES | YES | YES | 23128233 |
|  | 3 | rs11718165 | 2.04e-08 | YES | YES | YES | 21102463 |
|  | 5 | rs6596075 | 3.11e-06 | YES | YES |  |  |
|  | 5 | rs17234657 | 2.42e-12 | YES | YES | YES | 18587394 |
|  | 5 | rs11747270 | 1.05e-06 | YES | YES | YES | 18587394 |
|  | 7 | rs7807268 | 5.43e-06 | YES |  | YES | 26192919 |
|  | 10 | rs10883371 | 5.23e-08 | YES | YES | YES | 21102463 |
|  | 10 | rs10761659 | 1.69e-06 | YES | YES | YES | 22936669 |
|  | 16 | rs2076756 | 7.55e-15 | YES | YES | YES | 21102463 |
| Hypertension (HT) | 1 | rs10889923 | 1.38e-05 |  |  |  |  |
|  | 15 | rs2398162 | 6.01e-06 | YES |  |  |  |
| **Rheumatoid arthritis (RA)** | 1 | rs6679677 | <1.0e-15 | YES | YES | YES | 20453842 |
|  | 4 | rs3816587 | 7.28e-06 | YES | YES |  |  |
|  | 6 | rs9272346 | 7.38e-14 | YES | YES |  |  |
|  | 22 | rs743777 | 1.01e-06 | YES |  | YES | 23143596 |
| **Type 1 diabetes (T1D)** | 1 | rs6679677 | <1.0e-15 | YES | YES | YES | 19430480 |
|  | 2 | rs231726 | 1.43e-05 |  |  | YES | 30659077 |
|  | 4 | rs17388568 | 3.07e-06 | YES | YES | YES | 21829393 |
|  | 6 | rs9272346 | <1.0e-15 | YES | YES | YES | 18978792 |
|  | 12 | rs17696736 | 1.56e-14 | YES | YES | YES | 18978792 |
|  | 12 | rs11171739 | 8.36e-11 | YES |  | YES | 19430480 |
|  | 13 | rs4769283 | 1.20e-05 |  |  |  |  |
|  | 16 | rs12924729 | 7.86e-08 | YES | YES | YES | 17554260 |
| **Type 2 diabetes (T2D)** | 2 | rs6718526 | 1.00e-05 |  | YES | YES | 20418489 |
|  | 4 | rs1481279 | 9.44e-06 | YES | YES | YES | 28869590 |
|  | 6 | rs9465871 | 3.38e-07 | YES | YES | YES | 21490949 |
|  | 10 | rs4506565 | 5.01e-12 | YES | YES | YES | 23300278 |
|  | 12 | rs1495377 | 7.21e-06 | YES | YES | YES | 22885922 |
|  | 16 | rs7193144 | 4.15e-08 | YES | YES | YES | 22693455 |

The DeepCOMBI method finds 39 significant associations. According to the fifth column of **Table 2** 31 of these SNPs were also discovered by the traditional RPVT approach, because they have *p*-values <10^−5^. The other 8 of those 39 SNPs have *p*-values >10^−5^ and were hence not determined to be associated with the disease with RPVT in the original WTCCC publication. They are of special interest, because they represent additional SNP disease associations which the traditional analysis of the data was not able to identify. Out of these 8 novel discoveries, 6 have been validated independently in later GWAS or meta analyses: rs7570682 on chromosome 2 and rs1375144 on chromosome 2 for bipolar disorder; rs6907487 on chromosome 6 for coronary artery disease; rs12037606 on chromosome 1 for Crohn’s disease; rs231726 on chromosome 2 for type 1 diabetes and rs6718526 on chromosome 2 for type 2 diabetes.

On the other side, 2 out of the 8 novel DeepCOMBI SNPs with *p*-values > 10^−5^ have not yet been replicated in any independent GWAS or meta analyses. They have also not been identified by the COMBI method. Those entirely novel DeepCOMBI discoveries are rs10889923 on chromosome 1 for hypertension and rs4769283 on chromosome 13 for type 1 diabetes. To determine whether those two SNPs are biologically plausible discoveries for an association with the respective disease, their genomic regions were investigated in terms of functional indicators. Strong evidence of potential functional roles in the diseases were found. Firstly, rs10889923 maps on an intron for *NEGR1* (neuronal growth receptor 1), a very important gene many times linked to obesity, body mass index, triglycerides, cholesterol, etc. and many other phenotypes highly correlated with hypertension^66,67,68^. Even though *NEGR1* has been associated with many phenotypes in the GWAS Catalog, no GWAS has yet been able to directly link it to hypertension. Furthermore rs10889923 is part of a high LD region (according to LDmatrix Tool^69^) with variants that have been reported to be significantly associated with a number of psychiatric disorders and phenotypes, e.g. educational attainment (in Lee et al.^17^ rs12136092 with *p*-value < 1*e*^−11^ and a degree of LD *R*^2^ = 0.86 to rs10889923; rs11576565 with *p*-value < 1*e*^8^ and *R*^2^ = 0.63). This link suggests a potential connection between hypertension and related phenotypes with mental traits. rs10889923 can thus altogether be considered an excellent candidate for association with hypertension.

Secondly, rs4769283 on Chr. 13 lies in an intergenic region very close to a gene called MIPEP (Mitochondrial peptidase) which cannot be directly linked to T1D, but is reported as a significant eQTL (expression quantitative trait locus) for two other genes, namely C1QTNF9B and PCOTH^70^. Thus, MIPEP and therefore rs4769283 significantly control expression levels of mRNAs from these two genes in a particular tissue. Most remarkably, rs4769283 is a significant eQTL (with *p*-value = *1.1e* ^6^) for C1QTNF9B (Complement C1q And Tumor Necrosis Factor-Related Protein 9B) in (amongst several other tissues) the pancreas, which produces very little or no insulin in T1D patients. So even though the association of rs4769283 with Type 1 diabetes is not an obvious one, it is indeed an interesting novel discovery of the DeepCOMBI method.

To present a more condensed view of these discoveries Table 3 summarizes the findings of the three competitor methods, RPVT, COMBI and DeepCOMBI. When no screening step is conducted and RPVT *p*-values are calculated for all SNPs, 68 locations with *p* < 10^−5^ were identified as significant RPVT hits. COMBI and DeepCOMBI both apply a learning based SNP preselection step and thus, find fewer significant associations. The DNN based approach to this is seen to be more conservative than the SVM based one with only 39 identified locations of DeepCOMBI in comparison to 53 findings of the COMBI method. Even though the DeepCOMBI method finds fewer significant SNPs, the number of independently replicated SNPs of DeepCOMBI (= 31 replicated SNPs, yielding a precision of 79%) is identical to that of COMBI (31, precision = 58%) and almost identical to that of RPVT (33, precision = 49%). In addition, the DeepCOMBI method misclassified only 8 of all unreplicated SNPs as associated with the disease (yielding an error rate of only 21%), while RPVT wrongly classified 35 SNPs (error rate = 51%) and the COMBI method made 22 mistakes (error rate = 42%). These observations are quantified with pairwise one-sided Fisher’s exact tests for the null hypothesis of equal error rates for both methods. They produce significant *p*-values for both the comparison of DeepCOMBI vs. RPVT (Fisher’s exact test *p*-value of 0.00106) and the comparison of DeepCOMBI vs. COMBI (*p*-value = 0.01910).

**Table 3.** Quantitative summary of the significant findings of RPVT, COMBI and DeepCOMBI. For each of the three competitor methods the numbers of replicated and unreplicated hits (i.e. the number of true and false positives) as well as precision and error rates are presented. Pairwise tests for the null hypothesis of identical distributions for DeepCOMBI and the two baseline methods are performed and corresponding *p*-values given.

|  | Number of significant SNPs of |  |  |
| --- | --- | --- | --- |
|  | RPVT | DeepCOMBI method | COMBI method |
| **SNPs that have achieved $p < 10^{-5}$ in at least one external study** | 33 (49% precision) | 31 (79% precision) | 31 (58% precision) |
| **SNPs that have not achieved $p < 10^{-5}$ in an external study** | 35 (51% error rate) | 8 (21% error rate) | 22 (42% error rate) |
| **Overall** | 68 | 39 | 53 |
| **Pairwise $p$-value (one-sided Fisher's exact test)** | DeepCOMBI vs. RPVT = 0.00106<br>DeepCOMBI vs. COMBI = 0.01910 |  |  |

Instead of investigating the significant findings of the three competitor methods achieved by matching a specific error level, it is also possible to examine the performance of those methods for different levels of error. By increasing the significance threshold of each method from very conservative (*t*^*^ = 0, no significant SNPs) to very liberal (*t\** = ∞, all SNPs significant), we investigate here, how the three methods perform under these circumstances. In **Figure 6** we present the corresponding ROC and PR curves, where we interpret the replication of SNPs according to the GWAS catalog as a validation, i.e. we count a SNP as a true positive if it has achieved *p* < 10^−5^ in at least one external study. Overall, the findings obtained by the DeepCOMBI method are better replicated than those obtained by RPVT and COMBI for all levels of error. The performance metrics of the DeepCOMBI method (pink line) are consistently better than that of RPVT (light blue lines) and COMBI (dark blue lines). The DeepCOMBI method finds more true positives for different levels of error and yields higher levels of precision for different levels of recall than COMBI and RPVT. **Figure 6** also shows the performance curves of the other two baseline methods that threshold SNPs solely based on raw LRP relevance scores or raw SVM weights, respectively. As we can view these two methods and RPVT as the individual components of the combinatorial approaches and neither of these three can achieve the same level of performance as COMBI and DeepCOMBI, it can be deduced that all subparts are essential. Only the combination of the two parts of the DeepCOMBI method (NN with LRP explanation and statistical testing) can achieve the desired performance increase.

**Figure 6:**
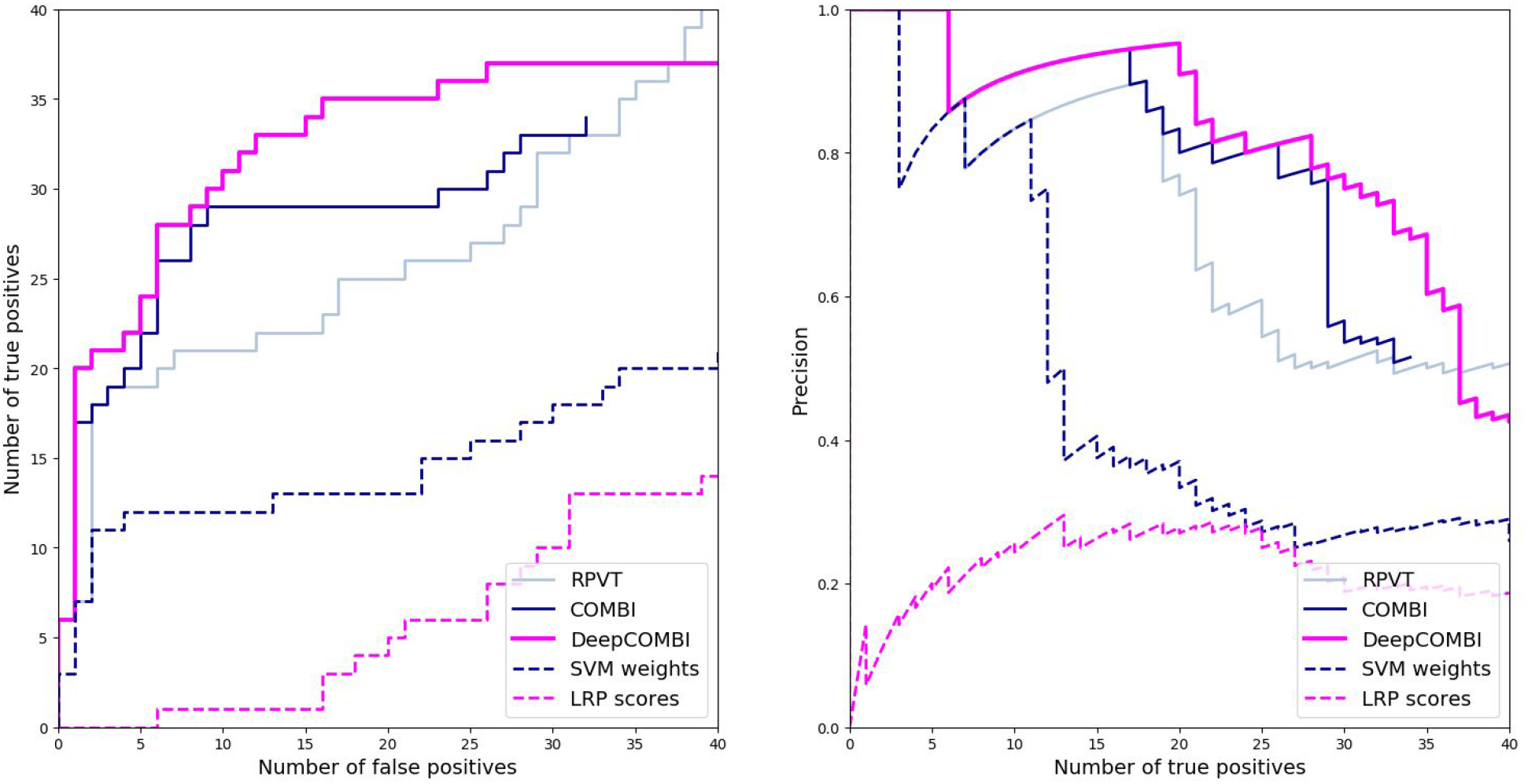
ROC and PR curves of DeepCOMBI and all competitor methods on WTCCC datasets. Performance curves of all methods averaged over all diseases and chromosomes are shown. ROC curves are presented on the left and PR curves on the right side. Replicability according to the GWAS catalog was used for validation.

### Concluding discussion

Numerous different approaches for the analysis of GWAS have been introduced since the first of its kind was published in 2002. Traditionally, they focus either on accurate phenotype prediction^1,2,3,4,5^ or the identification of SNP phenotype associations^6,7,8,9,10^. At first most of these approaches were of purely statistical nature^15,16^, but since machine learning has become increasingly important in data science, it has also found its way to the investigation of genetics data. A large range of all kinds of machine learning based tools have been proposed and investigated: regression and classification approaches, non-penalized and penalized methods, linear and nonlinear models^4,28,29,30,32,33^. A number of very well performing methods introduce the combination of traditional statistical testing concepts with more sophisticated machine learning tools^6,7,31^. With the increasingly larger amounts of available data, deep learning based approaches and artificial DNNs are now also being applied to GWAS datasets^41,42^. However, most of these publications focus on pure classification or regression prediction tasks^28,43,44,45^, rather than the identification of associated SNPs in the corresponding datasets^28,46^.

To fill this gap and firmly based on the combinatorial approach of the COMBI method^31^, the proposed DeepCOMBI method uses a deep learning based phenotype prediction in combination with statistical testing for the identification of SNPs that are associated with the phenotype under investigation. DeepCOMBI could be considered an extension of COMBI replacing the rather simple prediction tool of a linear SVM with a more sophisticated deep learning method and using the recent concept of explainability to uncover the decision making process of DNNs and extract SNP relevance scores via LRP^48,49,50^.

To our knowledge, Romagnoni et al.^28^ were the first and only scientists to use explainable AI in the context of GWAS and proposed to apply PFI. Even though they were able to identify some novel predictors, the prediction performance of their DNN was not better than that of traditional machine learning based tools. In addition, PFI is a generalized, model-agnostic approach and more sophisticated methods specifically tailored to DNNs are available. Hence, DeepCOMBI makes use of deep Taylor based explanation techniques by adopting LRP for the analysis of such GWAS data.

DeepCOMBI was shown to compare favorably to its main competitor COMBI on both generated controlled datasets as well as seven real-world GWAS datasets. These findings are in accordance with Romagnoni et al.^28^ who found that deep learning based methods can provide novel insights into the genetic architecture of specific traits. By applying LRP, we were able to leverage the power of DNNs and generate relevance scores that are less noise inflicted than the SVM importance scores of COMBI. In return, the pre-selection of candidates SNPs is better than that of COMBI and higher true positive rate and precision can be achieved for all levels of error. Since the COMBI method itself was shown before to outperform other combinatorial machine learning based approaches (Roshan et al.^6^, Meinshausen et al.^58^ and Wasserman and Roeder^59^) and a number of purely statistical analysis tools (Lippert et al.^21,27^), it can be directly deduced that DeepCOMBI also outperforms those approaches. For example, Wasserman and Roeder^59^ lose a great amount of statistical power by splitting the GWAS data under investigation into two parts performing SNP preselection on one part and statistical testing on the other. This approach is significantly less successful in identifying SNP disease associations than COMBI and hence DeepCOMBI who perform all substeps on the complete (and therefore statistically more powerful) dataset. Another exemplary statistical method which was shown to be outperformed by COMBI (and in consequence, DeepCOMBI) is based on linear mixed models (LMMs) proposed by Lippert et al.^21,27^. Even though they test for pairwise epistatic interactions in addition to the univariate tests and address the issue of population stratification in GWAS, they still test genetic locations and pairs thereof individually instead of simultaneously. In comparison to COMBI and DeepCOMBI which examine the genomic dataset as a whole, LMMs cannot achieve the same level of power and detection rates.

In addition to the main competitor method, COMBI, we also compared DeepCOMBI to the baseline methods of RPVT, raw LRP relevance scores and raw SVM importance scores and showed that only the combination of deep learning and multiple testing show the desired performance increase which cannot be achieved individually by one of these components.

A drawback of DeepCOMBI to consider might be that dense DNNs scale poorly with the number of SNPs studied, however, we have shown that DeepCOMBI performs well in combination with a *p*-value based SNP preselection step.

In conclusion, DeepCOMBI, a novel, AI based method was proposed for the analysis of GWAS data. After training a carefully designed DNN for the classification of subjects into their respective phenotype, the concept of explainable AI is applied by backpropagating the class prediction score to the input layer through the network via LRP. The resulting SNP relevance scores are used to select the most relevant SNPs for multiple testing in combination with a permutation based thresholding procedure. On both generated, controlled datasets as well as seven real GWAS datasets, DeepCOMBI was shown to perform better than a number of competitor methods in terms of classification accuracy of the DNN and in terms of ROC and PR curves when using either the generated labels or replicability in external studies as a validation criterion. In addition, two very promising, entirely novel SNP disease associations were discovered. Located on an intron for *NEGR1,* an important gene many times linked to obesity, body mass index and other correlated factors, rs10889923 on chromosome 1 was found to be significantly linked to hypertension. Another novel location found by DeepCOMBI to be associated with type 1 diabete**s**, is rs4769283. It is part of an intergenic region on chromosome 13 and was previously found to be an eQTL for C1QTNF9B in pancreas, the affected organ in T1D patients.

Future work on the subject of deep learning and explainable AI in the context of analyzing GWAS datasets could focus on one of the three steps of DeepCOMBI.

In the first step, DNNs with different architectures or other suitable analysis tools could be investigated. For example, future research could aim to harness the potential of convolutional networks^71^ in this application. By integrating multiple output nodes for multiple phenotypes, the DNN could also be extended to cover multivariate output variables and examine multimorbidities. DNNs can easily be adjusted to non-binary phenotypes.

Improvement ideas for the second step of the proposed method include the application of different explanation methods (e.g. PFI) or LRP backpropagation rules, for example according to the layer types, as advised by Montavan et al.^48^. Great potential lies in finding more sophisticated ways to combine the local LRP explanations of each individual subject to a single global explanation used for SNP selection. A very promising candidate would be a method called SpRAy^72^ which clusters the individual explanations and simplifies the identification of explanatory structures in subsets of subjects.

Future research work considering the third step of DeepCOMBI might examine the effects of replacing the χ^2^ test with a different, more sophisticated kind of test, e.g. investigating pairwise hypotheses or other multivariate effects.

## Acknowledgments

This work was supported by the German Federal Ministry for Education and Research through the Berlin Big Data Centre (01IS14013A), the Berlin Center for Machine Learning (01IS18037I) and the TraMeExCo project (01IS18056A). K.R.M. was supported in part by the Institute of Information & Communications Technology Planning & Evaluation (IITP) grant funded by the Korea Government (No. 2019-0-00079, Artificial Intelligence Graduate School Program, Korea University), and was partly supported by the German Ministry for Education and Research (BMBF) under Grants 01IS14013A-E, 01GQ1115, 01GQ0850, 01IS18025A, 031L0207D and 01IS18037A; the German Research Foundation (DFG) under Grant Math+, EXC 2046/1, Project ID 390685689. This work was also supported by Max Planck Society through the Max Planck School of Cognition. Correspondence to K.R.M.

## Author contributions

B.M., N.G., M. M.-C. H. and K.-R.M. designed and directed research; B.M., A.R. and J.R.F.H. performed research and analyzed data; and B.M., J.R.F.H., K.-R.M., and M. M.-C. H. wrote the paper.

The authors declare no conflict of interest.

